# AmyloGram Reveals Amyloidogenic Potential in Stroke Thrombus Proteomes

**DOI:** 10.1101/2025.07.07.663482

**Authors:** Douglas B Kell, Karen M. Doyle, J. Enrique Salcedo-Sora, Alakendu Sekhar, Melanie Walker, Etheresia Pretorius

**Affiliations:** Department of Biochemistry, Cell and Systems Biology, Institute of Systems, Molecular and Integrative Biology, University of Liverpool, Crown St, Liverpool L69 7ZB, UK; The Novo Nordisk Foundation Centre for Biosustainability, Building 220, Søltofts Plads 200, Technical University of Denmark, 2800 Kongens Lyngby, Denmark; Department of Physiological Sciences, Faculty of Science, Stellenbosch University, Stellenbosch Private Bag X1 Matieland, 7602, South Africa; Department of Physiology and Galway Neuroscience Centre, School of Pharmacy & Medical Sciences, University of Galway, Ireland; CÚRAM–SFI Research Centre for Medical Devices, University of Galway, Galway, Ireland; Department of Neurology, The Walton Centre, Lower Lane, Fazakerley, Liverpool L9 7LJ, UK; Department of Neurological Surgery, University of Washington, Seattle, WA 98104, USA

**Keywords:** amyloid, fibrinaloid, proteomics, inflammation, amyloidogenic sequences, stroke, cardioembolic, atherothrombotic

## Abstract

**Background:** Amyloidogenic proteins play a central role in a range of pathological conditions, yet their presence in thrombi has only recently been recognized. Whether computational prediction tools can identify amyloid-forming potential in thrombus proteomes remains unclear.

**Methods:** AmyloGram is a computational tool that estimates amyloid-forming potential based on n-gram sequence encoding and random forest classification. Using AmyloGram, we analyzed 204 proteins tagged by humans as amyloidogenic in UniProt. We then applied the same approach to proteins identified in thrombi retrieved using mechanical thrombectomy from patients with cardioembolic and atherothrombotic stroke. In addition we used AmyloGram to analyse the amyloidogenicity of 83,567 canonical human protein sequences.

**Results:** Among the UniProt-annotated ‘amyloid’ set, nearly all proteins received AmyloGram scores above 0.7, including 23 of the 24 human proteins. Even the lowest-scoring human protein, lysozyme (scoring 0.675), is known to form amyloid under certain conditions. In thrombi from both stroke subtypes, all detected proteins had AmyloGram scores above 0.7, suggesting a high likelihood of amyloid content. A majority of unannotated proteins also achieve AmyloGram scores exceeding 0.7.

**Conclusions:** AmyloGram reliably identifies known amyloid-forming proteins and reveals that stroke thrombi are enriched for proteins with high amyloidogenic potential. These findings support the hypothesis that thrombus formation in stroke involves amyloid-related mechanisms and warrant further investigation using histological and functional validation.

## INTRODUCTION

Proteins can adopt multiple stable conformations or macrostates. One important family of such conformations is represented by amyloids, which differ markedly from the native structures synthesized on the ribosome [1–12]. Amyloidogenic proteins, long studied in the context of neurodegeneration and systemic amyloidoses [3, 13–18], are increasingly recognized for their roles in vascular and thrombotic pathologies [19–25]. Recent evidence suggests that amyloid proteins may also contribute to clot formation and persistence in ischaemic stroke [26, 27]. Although ‘irreversibly’-formed amyloids (see [28]) are typically separated from native forms by a high energy barrier [29–33], they are often significantly more stable [29, 31, 34] and tend to form insoluble fibrils that can oligomerize. These properties are conferred by a defining structural feature, the cross-β motif [8, 35–39], which consists of β-sheets aligned perpendicular to the fibril axis. These sheets typically form stacks stabilized by hydrogen bonds and can assemble into protofibrils and eventually mature amyloid fibrils (Figure 1).

**Figure 1.**
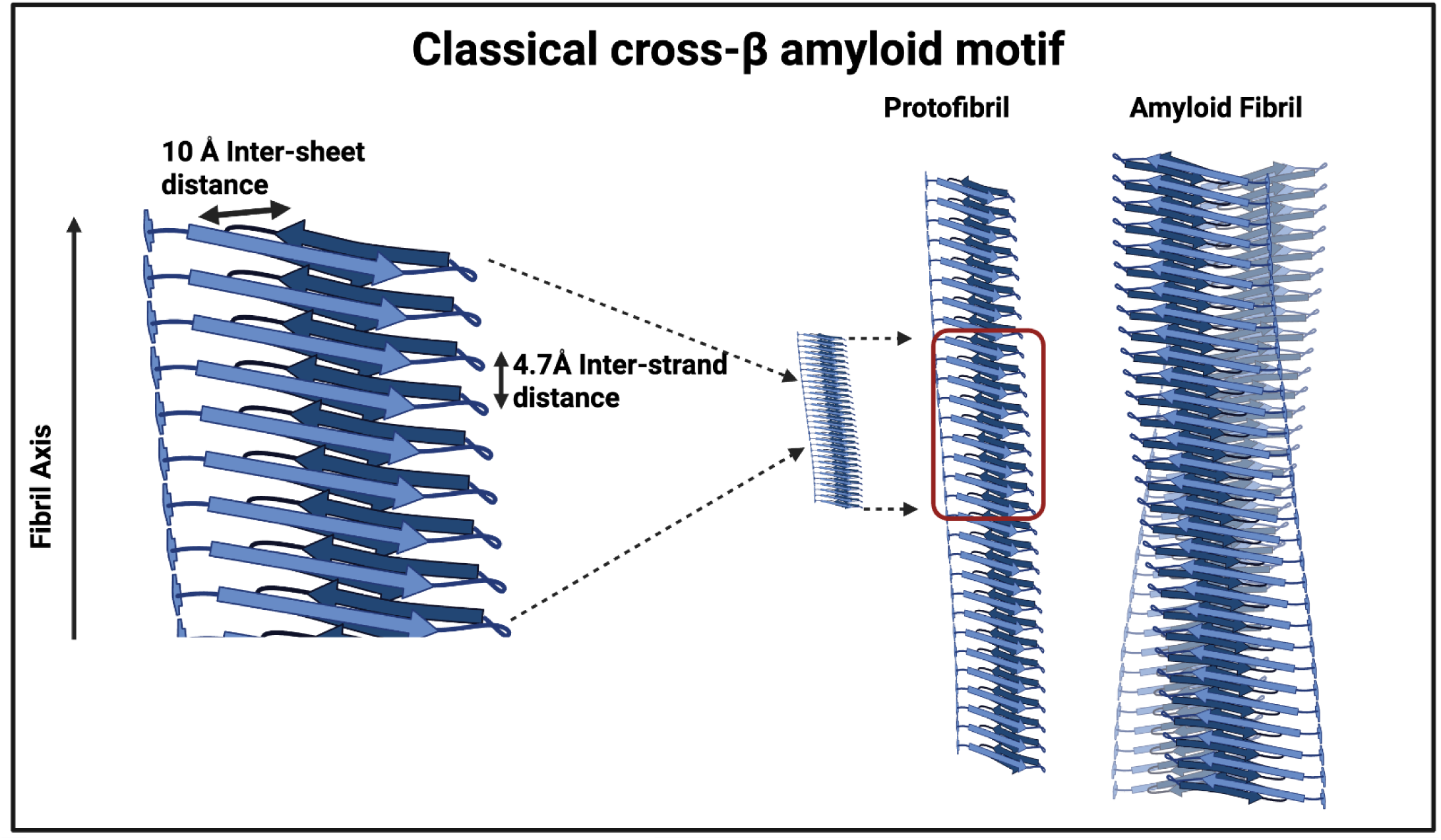
The basic cross-β-sheet structure underlying amyloid proteins. In each sheet, the β-strands (light and dark blue arrows) are aligned perpendicular to the fibrillar axis with a spacing of 4.7 Å and stabilized via hydrogen bonds. (Here these are antiparallel, but in other variants they are parallel.) The cross-β-sheet structure is formed by association of β-sheets at a distance of approximately 10 Å (it can vary from 6-12 Å depending on the sidechans involved [35, 40]). The further assembly of such cross-β-sheet structures results in the formation of protofibrils, and continued aggregation leads to the mature amyloid fibril. Adapted from a CC-BY 4.0 publication [41] using Biorender.com.

Almost any protein is capable of forming amyloid structures [40, 42], provided it contains a short run of amyloidogenic sequences [43, 44]. The SARS-CoV-2 spike protein provides a highly pertinent example [45–47], with its demonstrated ability both to form amyloid and to cross-seed amyloid formation [48], including during blood clotting [49, 50].

Given the diversity of sequences capable of forming amyloid, numerous computational tools (recently summarised [24]) have been developed to predict amyloidogenic potential based on sequence features. In this study, we focus on AmyloGram [51, 52], a predictive algorithm that uses n-gram encoding of amino acid sequences and a random forest [53] classifier to assign a single score between 0 and 1, reflecting overall amyloid-forming potential. In addition to a global score, the server provides residue-level predictions across the protein sequence, allowing for regional analysis of amyloidogenicity.

The present study aimed to assess computationally whether proteins identified in ischaemic stroke thrombi exhibit high levels of amyloid-forming potential consistent with the presence of amyloid as observed histologically, as previous work has demonstrated amyloid content in thrombi retrieved by mechanical thrombectomy from patients with acute ischaemic stroke [26, 27]. Based on these findings, we hypothesized that stroke thrombi are enriched in proteins with high intrinsic amyloidogenicity. To test this, we used AmyloGram as a validated computational prediction tool to assess amyloid potential across known amyloidogenic proteins and thrombus proteomes. The findings provide insight into possible molecular contributors to amyloid-driven clot formation and stability in the context of cerebrovascular disease.

## METHODS

### Amyloidogenicity Prediction Using AmyloGram

AmyloGram [51, 52] was used to estimate the amyloid-forming potential of protein sequences. This tool employs overlapping n-gram encoding of amino acid sequences and random forest classification to assign a global score ranging from 0 to 1, reflecting the likelihood of amyloidogenicity. It also produces residue-level profiles showing localized amyloidogenic potential across the protein sequence (some examples in [24] and below).

### Reference Dataset of Verified Amyloid Proteins

To calibrate the AmyloGram scoring framework we analyzed 204 proteins tagged as amyloidogenic in UniProt and marked as manually reviewed. (The specific query is https://www.uniprot.org/uniprotkb?query=%28keyword%3AKW-0034%29&facets=reviewed%3Atrue.) This dataset included entries from 95 species, with 24 annotated as human proteins. For all threshold-based analyses and clinical interpretation, we restricted comparative calibration and scoring to the 24 human proteins, given their greater biological and translational relevance. We recognise that some known amyloidogenic human proteins do not appear, but as our chief purpose was to establish the kinds of values in Amylogram that light up known amyloids our purposes are adequately served. Canonical full-length FASTA sequences were retrieved. Isoforms, fragments, and signal peptide trimming were excluded. Proteins were submitted in batches of approximately 40 to the AmyloGram web server http://biongram.biotech.uni.wroc.pl/AmyloGram/, and global scores were recorded and compiled for downstream comparison (Supplementary Table 1).

### Additional analysis using the R version of AmyloGram

The Uniprot version of the entire human proteome (39,675 kb) was downloaded from https://www.uniprot.org/uniprotkb?query=proteome:UP000005640 on 1/7/25, and consisted of 83,587 individual fasta sequences. These were analysed using the R version of AmyloGram https://github.com/michbur/AmyloGram. A very small subset of 20 proteins did not run (for reasons that were unclear), leaving a total of 83,567; since this represents 99.98% of them we do not consider that it will have any effect on the conclusions drawn. Default parameters were used for all analyses.

### Stroke Thrombus Proteomic Datasets

We applied the same scoring method to proteins identified in thrombi from patients with acute ischaemic stroke. Two mass spectrometry–based datasets were used. The first included 14 proteins found to be enriched in thrombi from patients with cardioembolic stroke [54]. The second included six proteins reported to be more abundant in thrombi from patients with atherothrombotic stroke compared to those with cardioembolic stroke [55]. Protein names or gene symbols were mapped to their corresponding UniProt accessions, and canonical human sequences were retrieved for analysis.

### Score Threshold and Comparative Analysis

AmyloGram scores were interpreted relative to the calibration set. A threshold of 0.7 was used to indicate high amyloid-forming potential, based on its consistent alignment with verified amyloidogenic proteins in the UniProt set. Proteins from stroke thrombi were evaluated individually, and their scores were compared across the two stroke subtypes to assess whether thrombus type was associated with predicted amyloidogenicity.

### Software and Data Management

Protein identifiers, sequence retrieval, and result collation were managed using Microsoft Excel. All UniProt IDs, AmyloGram scores, and relevant metadata are included in Supplementary Tables 1 and 2.

## RESULTS

### AmyloGram Scores Across Verified Amyloid Proteins

A total of 204 proteins were retrieved from UniProt using the “reviewed” plus “amyloid” keyword filters. These represented 95 species and included 24 human proteins. The most frequent entry was the major prion protein, which accounted for 70 of the 204 sequences. All 204 sequences were submitted to AmyloGram, and global amyloidogenicity scores were obtained. Five proteins received AmyloGram scores below 0.5. The remaining proteins showed a range of scores, with the great majority scoring above 0.7 (Figure 2).

**Figure 2.**
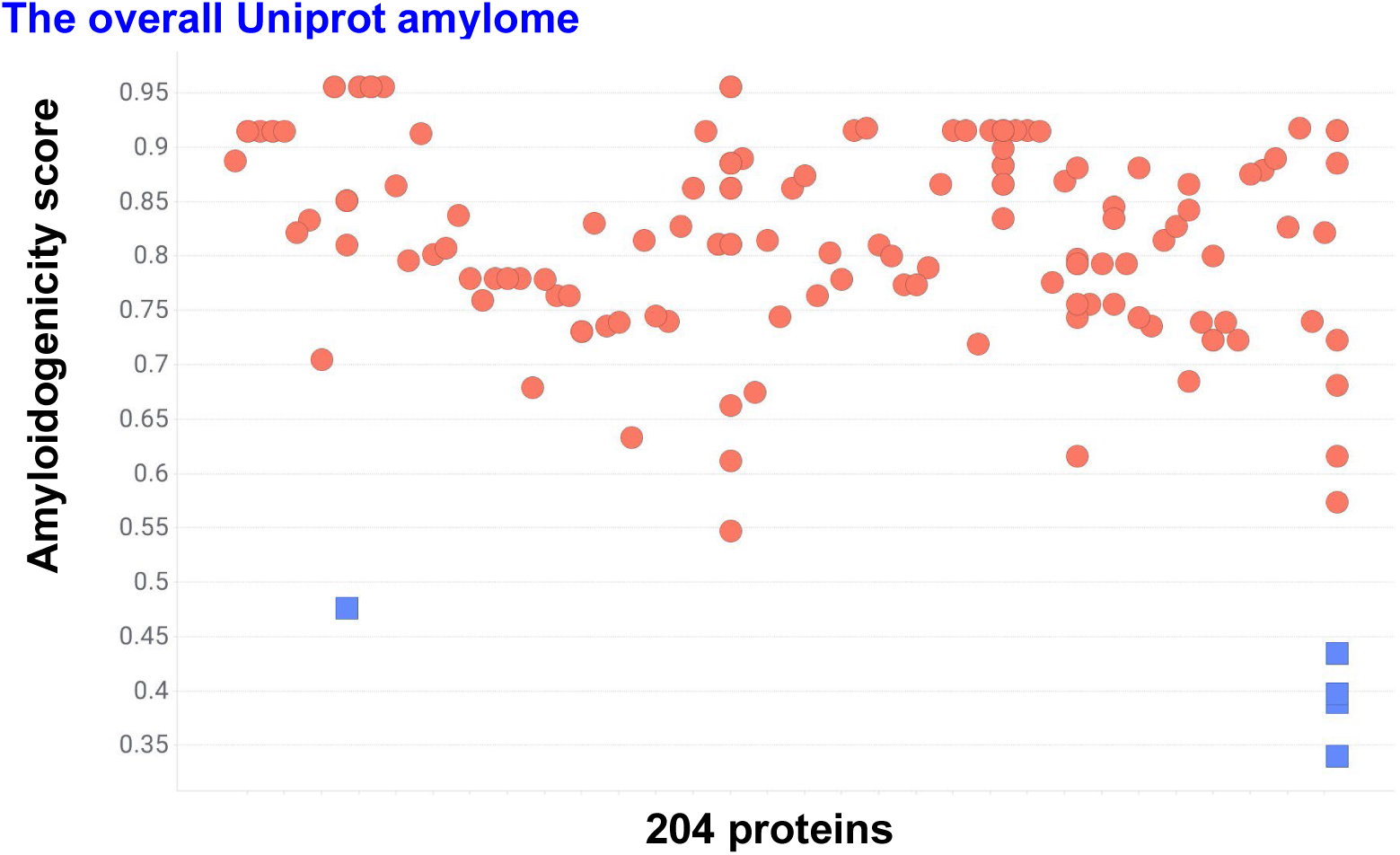
AmyloGram-predicted amyloidogenicity scores for 204 proteins annotated as amyloidogenic in UniProt. Each protein was assigned a global score between 0 and 1 based on sequence features. Proteins scoring below 0.5 indicated by blue squares. The majority of proteins scored above 0.7, consistent with experimentally verified amyloid-forming potential.

### AmyloGram Scores in Human Amyloid Proteins

Among the 24 human proteins in the dataset, AmyloGram scores ranged from 0.675 to above 0.95 (Figure 3). The highest scoring proteins included amyloid precursor protein (APP, part of whose sequence forms the highly amyloidogenic Aβ protein [56]) and islet amyloid polypeptide (IAPP or amylin [57–61]), both exceeding 0.95. Other high-scoring proteins included serum amyloid P [62–64], β2-microglobulin [30, 65–68], protein phosphatase 5, the major prion protein [69–71], and transforming growth factor β1 [72]. Given its known tendency to form amyloid blood clots [31, 73, 74] that are rather resistant to fibrinolysis [75, 76], fibrinogen A, with a score of 0.8276, is also of notable interest (and see below). Two serum amyloid A isoforms scored 0.7932 and 0.8541.

**Figure 3.**
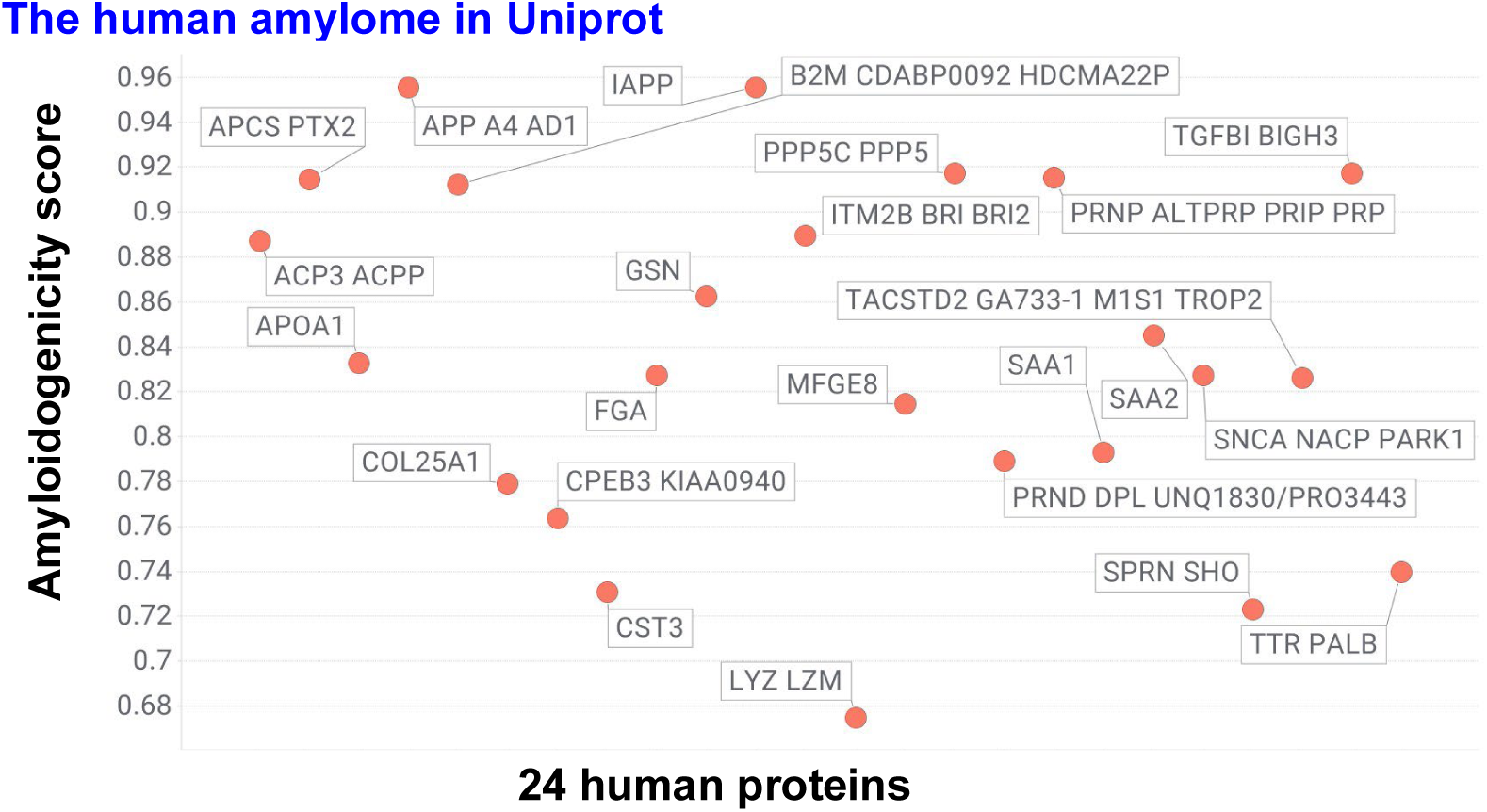
AmyloGram-predicted amyloidogenicity scores for 24 human proteins annotated as amyloidogenic in UniProt. Global scores are shown for each protein based on full-length canonical sequences. All proteins scored above 0.65, with several exceeding 0.9. Lysozyme, the lowest-scoring protein, still exceeded 0.67 and is known to form amyloid under specific conditions. Seven proteins scored above 0.9. Full names are given in Supplementary Table 1.

The four lowest-scoring human proteins tagged as amyloid were cystatin C, lysozyme, protein shadoo [77, 78], and transthyretin, with lysozyme scoring 0.675 and transthyretin scoring 0.74 (Figure 3). Consequently, we can safely conclude that any protein with a score above 0.7 is likely to be amyloidogenic, whether alone or by cross-seeding.

### Amylogram scores in the overall unannotated human proteome

Figure 4 show the distribution of the 83,567 human proteins at Uniprot for which Amylogram returned a score (Supplementary Table 2); 66,190 (79.2%) show a value exceeding 0.7 (Figure 4A), consistent with the view that ‘most’ proteins can indeed form amyloid under some circumstances. This observation is consistent with the hypothesis that a large fraction of human proteins contain local amyloidogenic segments and that sequence-based prediction tools such as AmyloGram are capturing real biochemical potential rather than overpredicting. There is also a minor tendency (Figure 4B, r^2^ ∼ 0.36 in lin-log coordinates, just 0.09 in lin-lin, neither line displayed) towards an increase in amyloidogenicity with length.

**Figure 4.**
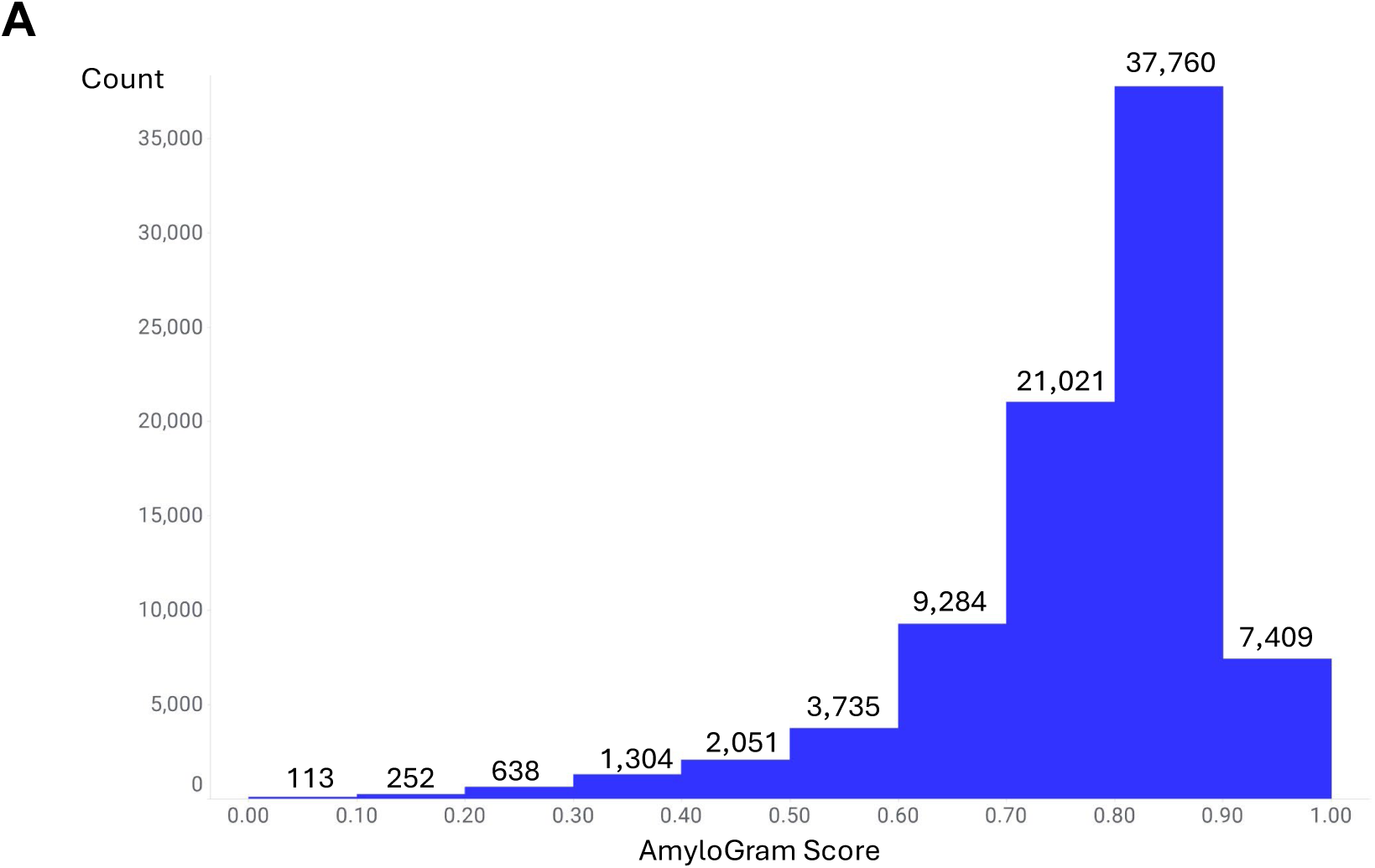
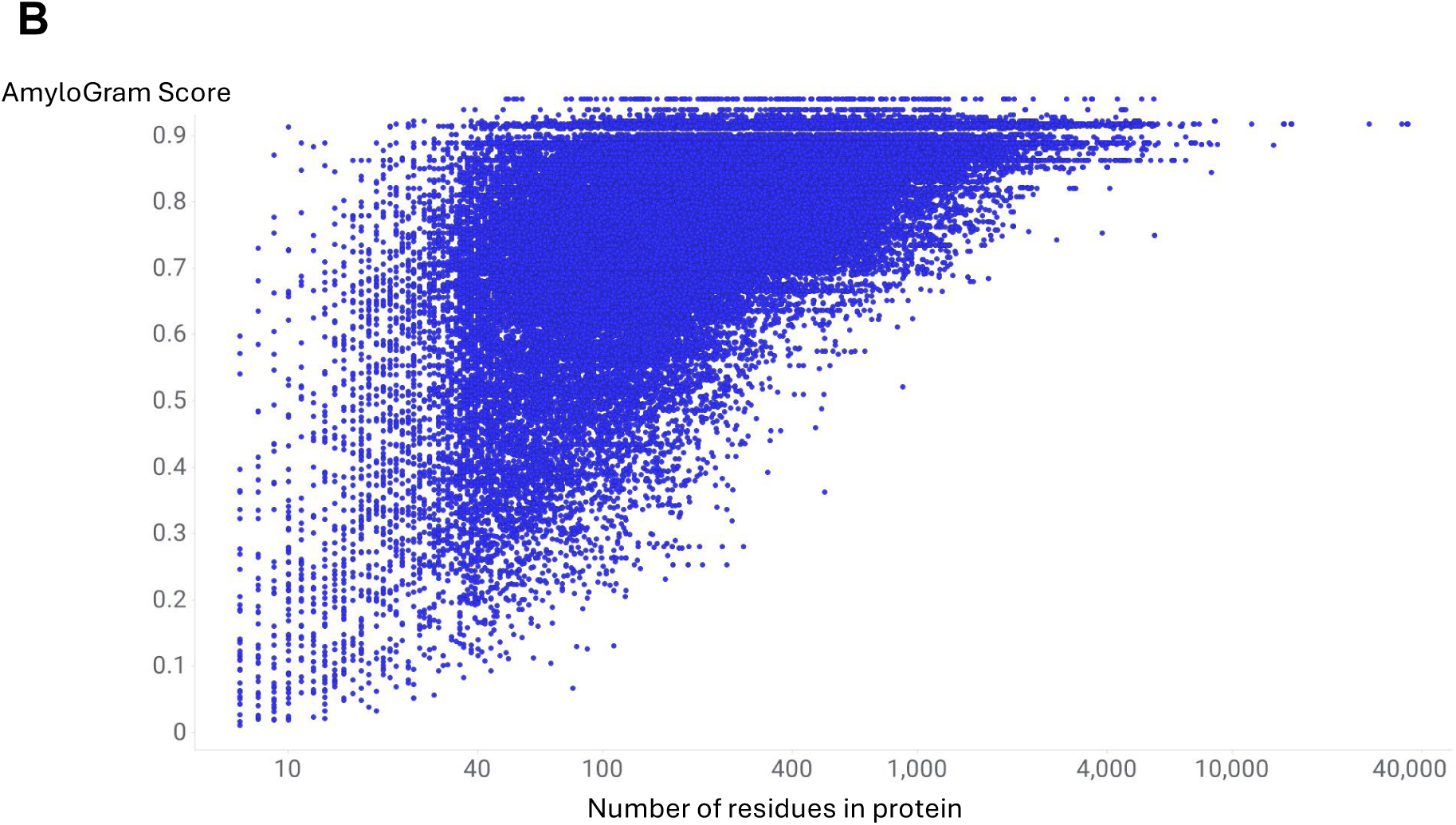
Distribution of AmyloGram scores across 83,567 full-length canonical human polypeptdies in UniProt. (A) Histogram of score frequency shows that 79.2% of sequences scored above 0.7, suggesting widespread presence of amyloidogenic motifs. (B) Scatter plot of score versus polypeptide length in amino acid residues shows a weak positive correlation, possibly reflecting increased opportunity for amyloid-prone motifs in longer proteins.Protein accession numbers and scores are listed in Supplementary Table 2.

### AmyloGram Scores in Stroke Thrombi by Subtype

We recently established that the thrombi removed by mechanical thrombectomy from patients who have suffered an ischaemic stroke also contain abundant amounts of amyloid material [26, 27]. In a previous strategy [25], we asked the question as to whether proteins additional to fibrinogen components that are known to occur in amyloid clots [24] can be used to predict the likely amyloid nature of those clots even when their amyloid status had not previously been measured. We here extend this strategy by asking the related question as to whether proteins observed experimentally (using proteomics) in stroke thrombi are also amyloidogenic <u>as judged by AmyloGram</u>. We analyzed proteins identified in thrombi from two published proteomic studies of ischaemic stroke. The first study reported 14 proteins enriched in thrombi from patients with cardioembolic stroke [54]. All 14 proteins received AmyloGram scores greater than 0.7 (Figure 5).

**Figure 5.**
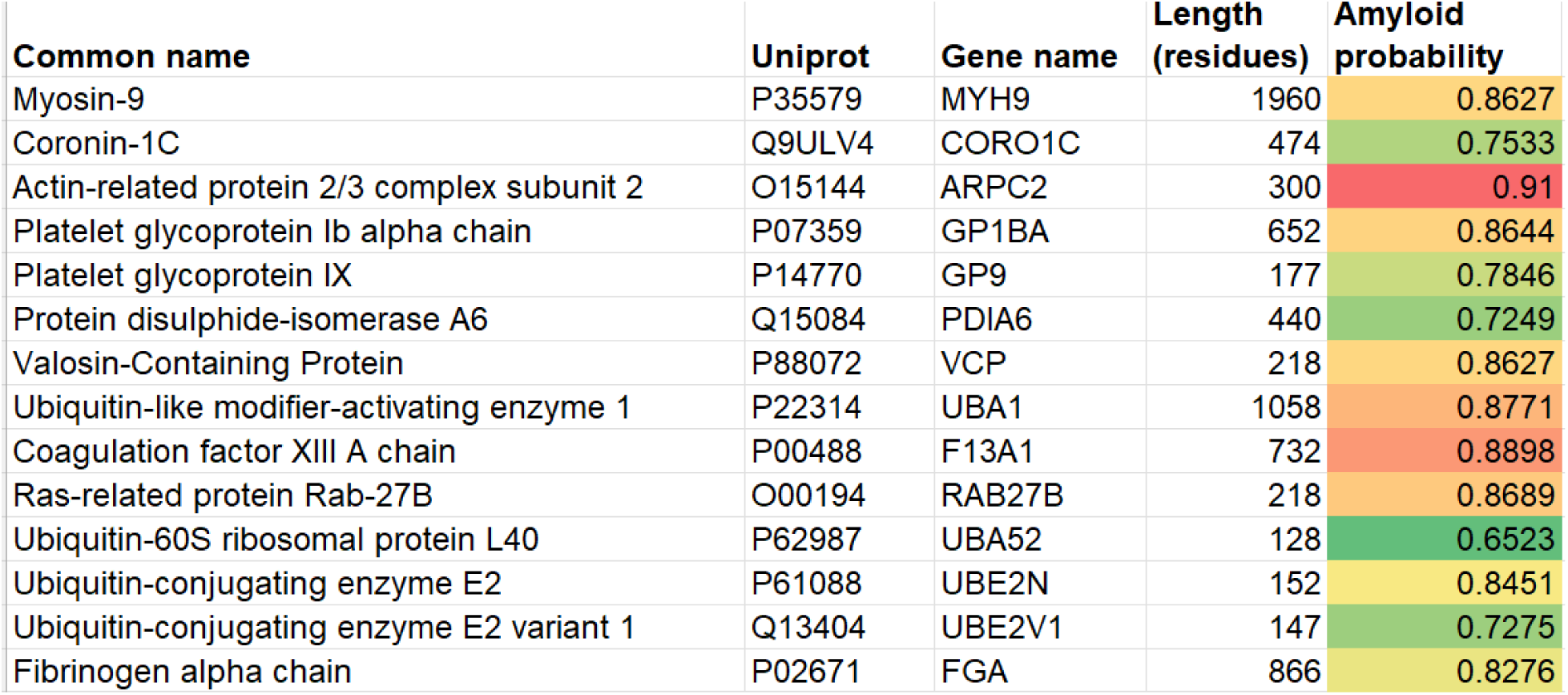
AmyloGram-predicted amyloidogenicity scores for 14 proteins identified in thrombi from patients with cardioembolic stroke, as reported in Table 1 of Rossi et al. [54]. Each protein was scored using its full-length sequence, and all received global scores above 0.7. Colour shading reflects the magnitude of the predicted amyloidogenicity.

The second study compared thrombi from cardioembolic and atherothrombotic stroke and identified six proteins with higher abundance in the atherothrombotic group [55]. These proteins were ATG3, RHD, FGA, SLC2A1, KRT1, and CLIC4. All six received AmyloGram scores greater than 0.75 (Figure 6).

**Figure 6.**
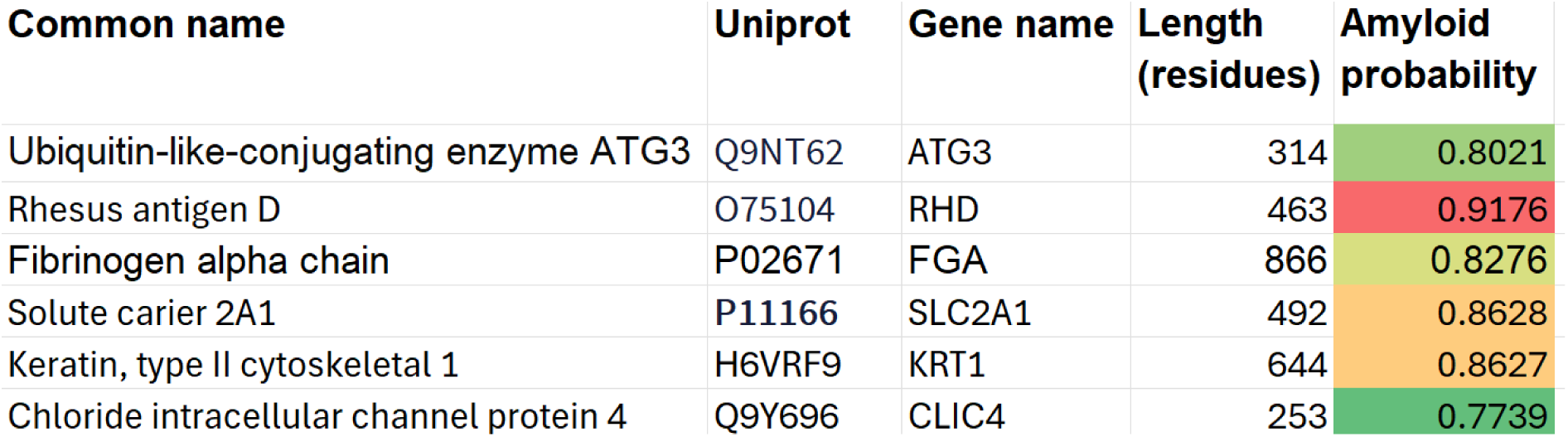
AmyloGram-predicted amyloidogenicity scores for six proteins reported to be more abundant in thrombi from patients with atherothrombotic stroke, based on the dataset from Lopez-Pedrera et al. [55]. All proteins received global scores above 0.75. Colour shading relates to the predicted amyloidogenicity score.

Although far from the most amyloidogenic of these proteins, a residue-level amyloidogenicity profile for ATG3 is shown in Figure 7.

**Figure 7.**
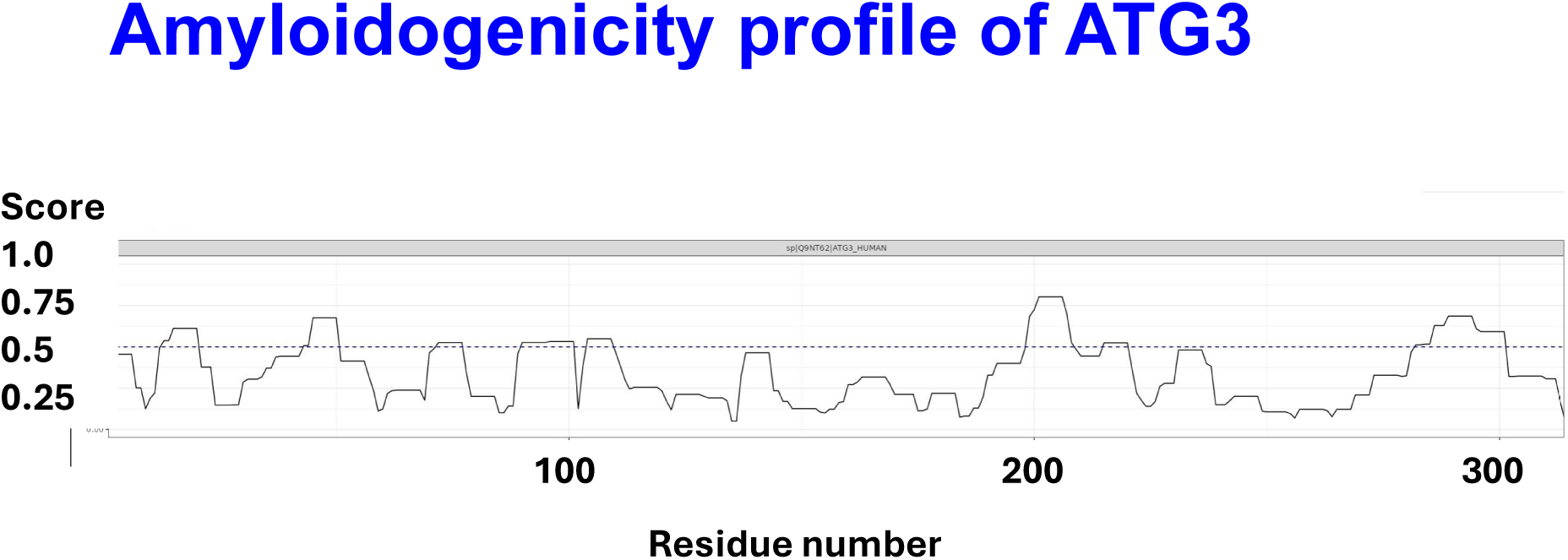
Residue-level amyloidogenicity profile of ATG3, generated using the AmyloGram server (accessed June 28, 2025). The plot displays predicted amyloidogenicity across the amino acid sequence, with several regions exceeding 0.5 and one run exceeding 0.75. This highlights localized segments with high fibril-forming potential within the full-length protein.

## DISCUSSION

Thrombi responsible for large vessel occlusion (LVO) in the setting of acute ischaemic stroke (AIS) are commonly characterized by a low recanalisation rate, even after intravenous thrombolysis (e.g. [79–88]). Regardless of their cause or variability, brain clots causing large vessel occlusion have an outer shell and core-shell structure that affects how easily they break down [89–91]. Studies using immunohistology have shown that these clots are quite diverse, with the thrombus outer shell being more resistant to tPA and thrombolysis generally than is the thrombus core [90]. Laboratory studies have also shown that clot shrinkage and compression can increase the clot’s resistance to breakdown [92–96], including by enhancing strong crosslinks between fibrin and other molecules. Unlike the fibrous fibrin in the inner core, the fibrin in the shell can appear dense and matted, and our previous work also demonstrated a tendency towards amyloidogenesis occuring in the shell of the clots examined [26, 27].

This study extends that work [26, 27] by evaluating whether computational prediction of amyloidogenicity aligns with experimentally verified protein behavior, and whether thrombi retrieved from patients with ischaemic stroke contain proteins predicted to be amyloid-forming. Using AmyloGram, we analyzed both a reference set of 204 human-reviewed amyloidogenic proteins from UniProt and protein lists derived from thrombus proteomic studies of cardioembolic and atherothrombotic stroke. In all cases, the majority of computationally assessed proteins, and each of those observed experimentally, scored above 0.7 at AmyloGram, consistent with their experimentally confirmed amyloid-forming capacity. The consistency of high amyloidogenicity scores across both stroke subtypes determined here implies that amyloid-related mechanisms may be a shared feature which drives the crosslinking between fibrin and other molecules, leading to a dense outer shell filled with tightly packed fibrin and platelets which are resistant to thrombolysis.

Among the 24 human proteins in the UniProt-verified dataset, scores ranged from 0.675 to greater than 0.95. Even lysozyme, with the lowest score of the 24 at 0.675, is well known to be capable of amyloidogenesis. Indeed, mutant forms of lysozyme are naturally amyloidogenic [97–99], and lysozymes are widely studied as an amyloidogenic model [98–106], albeit some require a rather acid pH to create stable amyloid conformations [107]. Cystatin C [108–111], with an AmyloGram score of just 0.73, and transthyretin [112–117] (0.74) are directly responsible for two well-known amyloidoses. These observations strongly support the use of a 0.7 threshold as a conservative predictor of amyloid-forming potential using AmyloGram.

We might have sought a ‘negative’ set of proteins that are not considered to form amyloid experimentally, but in fact this is not easily done. This is because it is considered that a majority of proteins can potentially form amyloid under suitable conditions [40, 42, 118, 119]; insulin is another well-known human example [120–126]), and the existence of a great many well-established amyloidogenic proteins above our nominal threshold of 0.7 is seen as sufficient justification for the approach. Note that AmyloGram does predict many proteins to have much lower amyloidogenicity (Figure 4), so we do not consider that it is pathologically over-predicting. Interestingly, even bacterial species produce proteins with strong amyloidogenic potential [127–131],where these structures may serve physiological roles such as structural defense or biofilm reinforcement[132].

The thrombus proteomic data demonstrated that every protein identified in both cardioembolic and atherothrombotic stroke clots received an AmyloGram score above 0.7. While proteomic profiling does not directly confirm the presence of amyloid structures (although it is consistent with a noticeable resistance to the kinds of protease commonly used in proteomics [75, 76]), this finding suggests that the protein composition of these thrombi is strongly biased toward sequences with intrinsic amyloidogenic features. These results align with prior histological evidence of amyloid deposition in stroke thrombi, as detected by thioflavin T [26, 27], and provides a straightforward explanation for the comparative resistance [133, 134] of these clots towards fibrinolytics such as tissue plasmin(ogen activator).

The consistency of high amyloidogenicity scores across both stroke subtypes implies that amyloid-related mechanisms may be a shared feature of thrombus formation or stabilization. The complex proteomes observed in these (and other [23, 75, 76]) clots are only realistically accounted for by a rather widespread cross-seeding [24, 25]. However, subtle differences likely exist, since the observed amyloid dispositions were highly heterogeneous both within and between clots [26, 27]. For example, fibrinogen A and serum amyloid A are known to co-aggregate [135] and were among the high-scoring proteins observed in the thrombus datasets. Future work could assess whether amyloid burden or composition varies by stroke mechanism, infarct size, or thrombus resistance to lysis.

Finally, this work raises the possibility that cross-seeding interactions between fibrin-based clot components and classical amyloid-forming proteins, such as APP, Aβ or IAPP, may contribute to the persistence or remodeling of thrombi. Such interactions have been observed in other settings [23, 136–141] and may link clot composition to post-stroke inflammation or neurodegeneration.

The residue-level profiles from AmyloGram, as illustrated for ATG3 (Figure 6), also suggest that specific regions within proteins may disproportionately contribute to amyloidogenesis. These localized high-scoring segments (‘hotspots’) could represent therapeutic or diagnostic targets, particularly if they contribute to fibril nucleation or structural stability.

Our findings demonstrate that proteins identified in ischaemic stroke thrombi consistently score highly for predicted amyloidogenicity, regardless of stroke subtype. This supports the experimental observation that amyloid-forming potential may contribute to thrombus structure or stability *in vivo*. The application of sequence-based tools such as AmyloGram provides a complementary strategy for identifying these features, especially in cases where traditional proteomic or histological methods may not detect underlying amyloidogenic signatures. These results motivate further investigation into the structural roles of amyloid-prone proteins in thrombotic disease and may inform future therapeutic or diagnostic approaches.

## ACKNOWLEDGEMENTS

This work originated from a collaborative discussion among the authors and was further developed through interdisciplinary exchange across the contributing groups.

## AUTHOR CONTRIBUTIONS

All authors contributed to the conceptualization of the study, data analysis, manuscript drafting, and final editing. All authors have read and approved the final version of the manuscript.

## DATA AVAILABILITY

All data supporting the findings of this study are available within the article and its Supplementary Materials. The UniProt human protein dataset used for calibration is publicly accessible at https://www.uniprot.org/uniprotkb?query=%28keyword%3AKW-0034%29&facets=reviewed%3Atrue. Stroke thrombus proteomic datasets were obtained from previously published studies [54, 55] and are cited accordingly. Amyloidogenicity scores for all analyzed proteins, as generated by the online AmyloGram server (http://biongram.biotech.uni.wroc.pl/AmyloGram/), are provided in Supplementary Table 1, and those from the R program in Supplementary Table 2.

## FUNDING

D.B.K. acknowledges support from the Balvi Foundation (grant 18) and the Novo Nordisk Foundation (grant NNF20CC0035580). E.P. received funding from the National Research Foundation of South Africa (grant 142142), the South African Medical Research Council (Self-Initiated Research grant), and the Balvi Foundation. The content and findings presented here are the sole responsibility of the authors and do not necessarily reflect the views of the funding agencies.

## CONFLICTS OF INTEREST

E.P. is a named inventor on a patent application related to the use of fluorescence-based methods for microclot detection in Long COVID. The funding bodies had no involvement in the design of the study, data collection or analysis, manuscript preparation, or the decision to submit for publication. The other authors have no disclosures.

